# Juvenile rank acquisition influences fitness independent of adult rank

**DOI:** 10.1101/812362

**Authors:** Eli D Strauss, Daizaburo Shizuka, Kay E Holekamp

## Abstract

Social rank has been identified as a significant determinant of fitness in a variety of species. The importance of social rank suggests that the process by which juveniles come to establish their position in the social hierarchy is a critical component of social development. Here, we use the highly predictable process of rank acquisition in spotted hyenas to study the consequences of variation in rank acquisition in early life. In spotted hyenas, rank is ‘inherited’ through a learning process called ‘maternal rank inheritance.’ This pattern is highly predictable: ~80% of juveniles acquire the exact rank predicted by the rules of maternal rank inheritance. This predictable nature of rank acquisition in these societies allows the process of rank acquisition to be studied independently from the ultimate rank that each juvenile attains. In this study, we use a novel application of the Elo-rating method to calculate each juvenile’s deviation from expected pattern of maternal rank inheritance during development. Despite variability in rank acquisition in juveniles, most of these juveniles come to attain the exact rank expected of them according to the rules of maternal rank inheritance. Nevertheless, we find that transient variation in rank acquisition in early life predicts long term fitness consequences for these individuals: juveniles ‘underperforming’ their expected ranks show reduced survival and lower lifetime reproductive success than better-performing peers. Finally, we present evidence that this variability in rank acquisition in early life represents a source of early life adversity, and that multiple sources of early life adversity have cumulative, but not compounding, effects on fitness.

## Introduction

Group living comes with both benefits and costs. Benefits such as reduced predation risk, cooperative breeding, and cooperative resource defense, are weighed against costs such as increased competition over local resources, pathogen transmission, and risk of social conflict. These costs and benefits may not be experienced by all group members equally; some individuals gain more of the benefits and suffer fewer of the costs than others [1,2]. In many animal societies, this disparity among group-mates is reflected by a dominance hierarchy, where individuals differ systematically in their tendency to display subordinate signals to their group-mates [3]. A useful abstraction of the network of complex and unequal relationships among group members is ‘rank,’ which describes the extent to which an individual is able to exert power over its group-mates. Extensive research from a variety of organisms has demonstrated that individuals of high rank, which are able to exert power over most other individuals in their social group, enjoy dramatic advantages as a result of their position in the social hierarchy, although species vary in the nature and strength of the relationship between social status and fitness [2,4–6].

Decades of work has demonstrated various correlates with dominance rank or status within a social group. For example in many species, the social ranks of adults are well predicted by certain phenotypes such as body size or physical markings, or certain conventions such as age or tenure [7–12]. Social factors, such as support from conspecifics or presence of kin, also influence dominance rank [5,13–15]. Winner- and loser-effects, where individuals that win (lose) a particular interaction show increased probabilities of winning (losing) subsequent interactions, have also been demonstrated to affect hierarchy formation in a number of species [16,17]. In many cases, the effects of these factors on rank is relatively strong such that one can predict the rank of an adult based on their phenotypes, demography, or ranks of relatives.

Although a vast literature now addresses the correlates of dominance ranks in groups, comparatively little is known about the processes governing rank acquisition, how individuals may experience variations in such processes, and how deviation from predicted dominance relations during development may affect future fitness. The process of social rank acquisition in juveniles may be highly complex and difficult to predict [15,18], as juveniles continually re-negotiate dominance relationships with their group-mates as they mature [19,20]. Yet, this process may have disproportionately large effects on later survival or reproduction, particularly in species that live in cohesive social groups throughout life, where the transition between juvenile social development and adult social behavior is gradual. Although signatures of early-life social networks have been shown to last into adulthood in some species [21–23], it is unclear whether dominance-related behaviors in early life have effects beyond influencing the ranks the juveniles ultimately attain as adults.

There are multiple reasons why the process of rank acquisition might relate to fitness, independent of the ranks juveniles ultimately acquire. First, social uncertainty is costly [24,25], and a tumultuous process of rank acquisition could be a source of significant social uncertainty, and thus adversity, in early life. Early-life adversity is associated with downstream consequences in many species [26–28], so costs of social uncertainty in early life could potentially have far reaching fitness consequences. Second, it is possible that fitness-associated phenotypes relate to the rank-acquisition process independently of the ranks individuals ultimately acquire, and thus variation in these phenotypes may influence both the rank acquisition process and fitness, but not rank in adulthood. Finally, adults may remember the outcomes of social interactions as juveniles, and thus early-life interactions might influence social relationships in adulthood independent of the rank the juvenile ultimately attains.

Here we take advantage of the social system of the spotted hyena (*Crocuta crocuta*) to conduct a large-scale prospective study on the consequences of variation in rank acquisition among juveniles. Spotted hyenas acquire their rank through a learning process known as maternal rank ‘inheritance’ with youngest ascendency. In this system, juveniles come to acquire the rank directly below that of their mothers and above those of their older siblings; this system is found in many Cercopithecine primates as well as in spotted hyenas. Prior work found that rank acquisition by this process is highly predictable: most (78.1%) females acquired the exact ranks predicted by maternal rank inheritance with youngest ascendency [13], and were consistently able to dominate lower-born adult females when they were around 18 months old [29]. Here, we show that there is considerable variation in the process of rank acquisition, independent of the rank the juvenile ultimately acquires. To measure variation in rank acquisition, we develop the ‘Elo-deviance’ method, which measures deviation from a hypothesized rank for each juvenile based on the rank of its mother relative to those of other adult females in her social group. We then relate Elo-deviance during development to survival and lifetime reproductive success, and find that this variability in rank acquisition has important fitness consequences, independent of the rank the juvenile ultimately acquires.

### A novel method to measure variation in rank acquisition

We developed a novel ‘Elo-deviance’ method to measure variation in rank acquisition among juveniles. The Elo-deviance method assesses deviation from an expected pattern of contest outcomes by calculating the difference between the observed Elo-rating for a focal individual and the Elo-rating that individual would have attained under some prior hypothesis. This method is based on the widely used Elo-rating method, which calculates a numerical dominance score for each individual in a social group by updating the relative dominance scores of individuals after each observed interaction [30,31]. Scores for the winner and loser of each interaction change in proportion to the expected probability of the observed outcome, as determined by their score prior to the interaction; expected outcomes lead to smaller changes in scores, whereas unexpected outcomes lead to larger changes. Thus, the Elo-rating method is more sensitive to unexpected outcomes than to expected outcomes.

In this study, the prior hypothesis we used in the Elo-deviance method is that of maternal rank inheritance, where the ranks among juveniles should be isomorphic with the ranks among their mothers. Thus, we calculate a juvenile’s Elo-deviance score by subtracting its observed Elo-rating from the Elo-rating it would have received had it won or lost every interaction as expected based on its mother’s social rank. Observed and expected Elo-ratings were calculated using the *aniDom* R package [32].

To ensure that any differences between an individual’s observed and expected Elo-rating is due to its behavior and not to the behavior of its group-mates, Elo-deviance scores are calculated for each individual independently. Thus, aggressive interactions are first restricted such that they involve only the focal individual, and interactions can be further restricted based on the study question (e.g., only interactions among members of the same sex, only interactions during a specific time period). Observed Elo-ratings are then calculated based on the observed outcomes of interactions; expected Elo-ratings are calculated from the same set of interactions with the outcomes determined according to the hypothesis under investigation. An Elo-deviance trajectory is calculated for the focal individual by subtracting its expected Elo-rating from its observed Elo-rating, and the Elo-deviance is determined as the difference between observed and expected Elo-rating after the final interaction. Individuals who win and lose interactions according to the hypothesis earn Elo-deviances close to 0, whereas individuals who lose unexpectedly or win unexpectedly earn Elo-deviances below or above 0, respectively. Numbers of points gained/lost are scaled according to a constant, K, which we set to 20 for this analysis (following [33]). We also ran the same analyses with K = 100 (following [30]) and this did not change the conclusions of the study (see Supplemental Materials).

To measure individual variation in rank acquisition, we assessed Elo-deviance for each juvenile at the end of their den-dependent period. Spotted hyenas spend most of the first year of their life at the communal den, where juveniles from multiple mothers within the group are raised together. This period is one of intense social development for these juveniles, and by the end of the den-dependent period, juvenile ranks within their den cohorts typically match the relative ranks of their mothers (their maternal ranks) [34]. Because juvenile’s acquire their ranks relative to their peers before developing relationships with the rest of their group-mates [29,34], we assessed Elo-deviance based on interactions with peers only. See Supplemental Materials for analyses of Elo-deviance in later life-history stages.

## Methods

### Field data collection

We examined the relationship between juvenile rank acquisition and fitness in spotted hyenas from four study groups (‘clans’) in the Maasai Mara National Reserve in south-west Kenya. Spotted hyenas live in large mixed-sex clans characterized by highly fluid fission-fusion dynamics [35], meaning that individuals from the same clan associate in subgroups that change composition several times per day. Demographic data were collected during daily morning and evening observation sessions between 1988 and 2019 for one clan and between 2008 and 2019 in three others. Aggressive interactions among individuals of all age classes were collected using all-occurrence sampling [36]; aggressive interactions were collected up until June 2016 for two clans, December 2016 for one clan, and March 2017 for the fourth clan. We used the aggressive interactions among adult females to infer maternal ranks (i.e., rank of a juvenile’s mother relative to other mothers) as in [13,37]; we used the aggressive interactions among juveniles to measure variation in rank acquisition using the Elo-deviance method. In all cases we used, only aggressive interactions in which the recipient displayed submissive behavior.

### Modeling survival

We modeled survival as a function of Elo-deviance at den independence using mixed effects cox proportional hazards models (using *coxme* R package [38]). Mortality was determined to have occurred when an individual was found dead or when at least 6 months passed without it being observed. Survival data were right-censored for all individuals who were still alive at the end of June, 2019. Among males, we were unable to distinguish unobserved mortality from dispersal after 2 years of age, so male mortality data were right-censored at 2 years old.

In addition to Elo-deviance, we also included maternal rank (calculated as the rank the juvenile’s mother held in the year of the juvenile’s birth), and coded it categorically as ‘high’ and ‘low’ rank based on they were in the upper or lower half of the hierarchy. Rank relationships among females were inferred yearly for all adult females who were at least 1.5 years old at the start of the calendar year using the Informed MatReorder method, as in previous studies [13,37,39]. To control for the possible influence of variable sampling on Elo-deviance measures, we included the number of interactions used to calculate Elo-deviance as a predictor in each model. Additionally, we included a binary predictor coding whether the juvenile’s mother survived until the juvenile reached adulthood (2 years old). Finally, we included a random effect of clan to account for variation at the the clan level.

Elo-deviance in all models was coded as a categorical predictor with two categories: Elo ≥ expected (i.e., Elo-deviance ≥ 0) and Elo < expected (i.e., Elo-deviance < 0). Models with Elo-deviance as a categorical predictor performed better than the same models with Elo-deviance as a continuous predictor (∆AIC = 5.207), with the raw Elo score (i.e., observed Elo score rather than Elo-deviance) as either a categorical predictor (Above/below expected; ∆AIC = 7.987) or a continuous predictor (∆AIC = 7.842), or a null model including other covariates but no measure of the state of rank acquisition at den independence (∆AIC = 6.225).

### Modeling lifetime reproductive success

We used Poisson generalized linear mixed effect models to assess the effects of Elo-deviance at den independence on lifetime reproductive success (LRS). LRS was calculated for the subset of the juveniles that were female and that died during the study (n = 147). We could not assess LRS for males because they dispersed and because we could rarely assign paternity to them. LRS was calculated as the number of offspring surviving to adulthood (2 years old) produced by each female. We included the same predictors in our models of LRS as we included in the survival analysis. Models were created using the *lme4* R package [40].

## Results

### General patterns of rank acquisition

Importantly, although Elo-deviance at den independence showed considerable variability (Figure 1a), most juveniles ultimately acquired their rank as predicted by maternal rank inheritance with youngest ascendency, regardless of their Elo-deviance at den independence (Figure 1b). Rank at the onset of adulthood was highly correlated with the mother’s rank in that year (Pearson’s r = 0.980; 95% CI = [0.971, 0.987]; n = 102), and 77.5% of new adults acquired their rank exactly according to maternal rank inheritance with youngest ascendency. A Chi-squared test revealed that Elo-deviance at den independence (Elo ≥ expected or Elo < expected) did not predict whether juveniles acquired a rank above expected, below expected, or exactly as expected according to maternal rank inheritance with youngest ascendency (χ-squared = 1.715, df = 2, p = 0.424).

**Figure 1.**
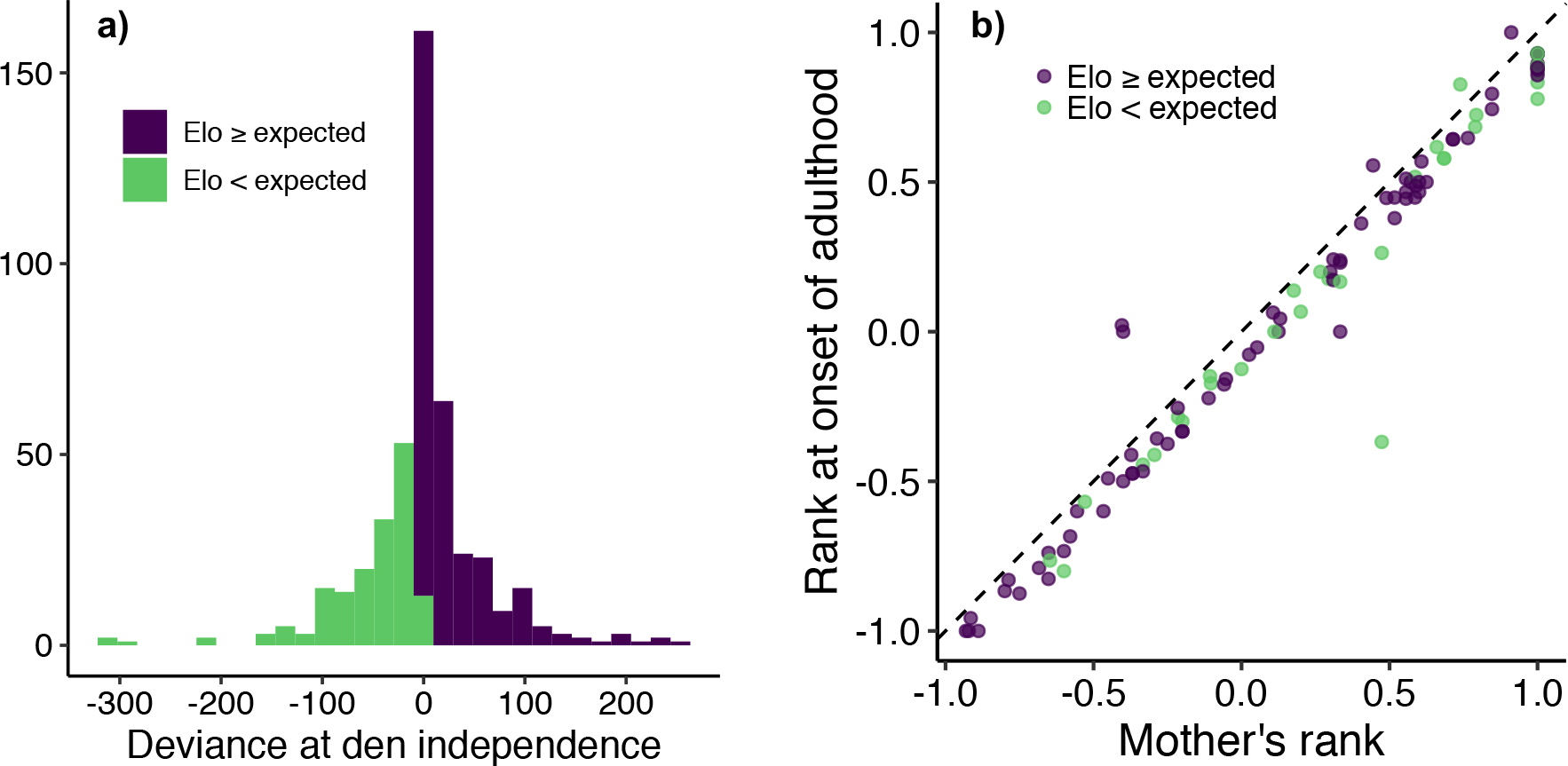
(a) Histogram of Elo-deviance at den independence. (b) The relationship between the juvenile’s mother’s rank and the juvenile’s rank at the onset of adulthood. According to maternal rank inheritance, points should lie directly below the dashed line (denoting where mother’s rank and juvenile’s rank are exactly equal). In this study, 77.5% of juveniles acquired the exact rank predicted by maternal rank inheritance. Elo-deviance at den independence (color) did not affect the rank attained by the onset of adulthood. Taken together, these plots show transient variability in rank acquisition at the end of the den-dependent life-history stage that fails to manifest in rank differences in adulthood.

### Fitness correlates of Elo-deviance at den independence

Elo-deviance at den independence significantly predicted survival (n = 465; Figure 2): Juveniles with deviance scores below 0 at den independence die earlier (hazard ratio = 1.529; 95% CI = [1.148, 2.037]; p = 0.004). Death of the juvenile’s mother prior to reaching adulthood (hazard ratio = 1.724; 95% CI = [1.255, 2.369]; p = 0.0008) also predicted reduced survival, but being of low maternal rank did not (1.215; 95% CI = [0.911, 1.620]; p = 0.180). All results reported here were from the full model, and thus control for the effects of the other predictors.

**Figure 2.**
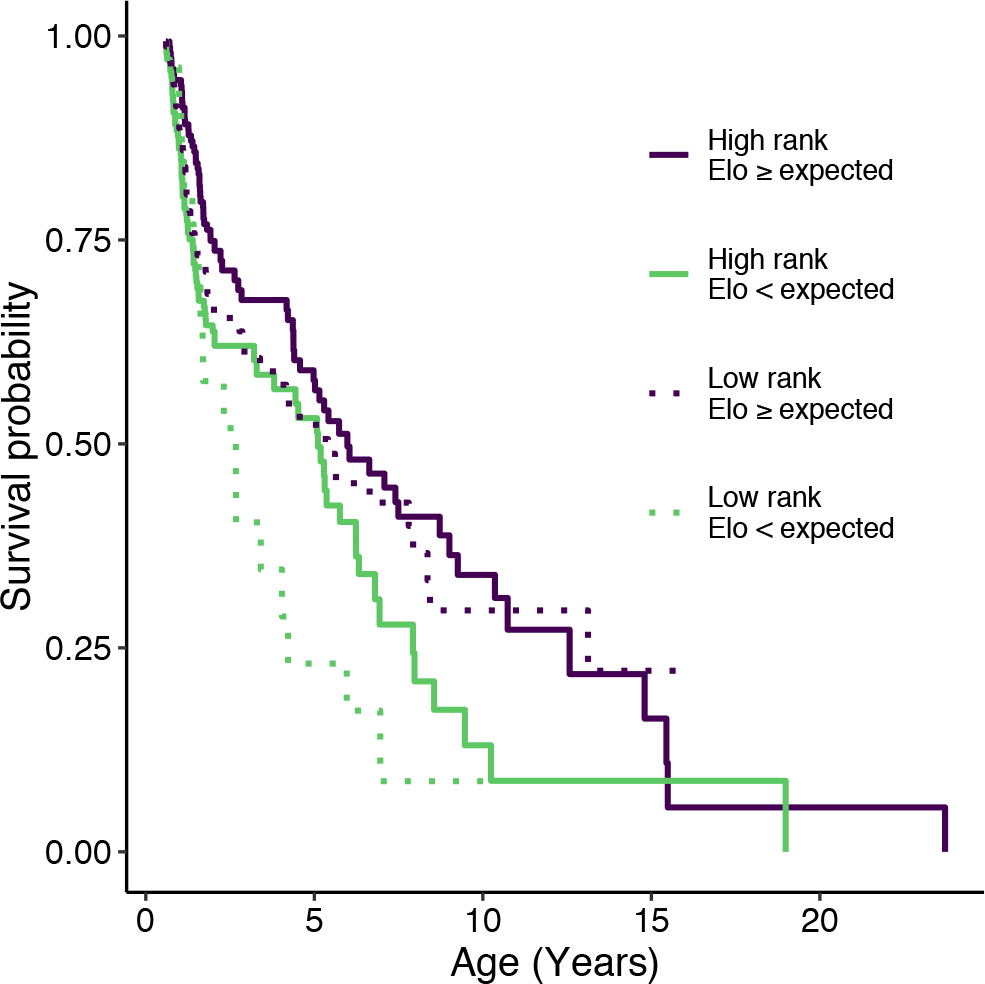
Survival probability as a function of Elo-deviance at den independence and maternal rank. Juveniles with below Elo-deviance < 0 showed reduced survival. Death of the mother before the juvenile reached adulthood also predicted reduced survival, but being born to a low-ranking mother did not predict survival after controlling for the other variables in the model.

Elo-deviance at den independence also predicted LRS (Figure 3); females with deviance scores below 0 at den independence produced fewer offspring than did females with deviance scores ≥ 0 (ß_Elo-deviance below 0_ = −0.490 ± 0.168, p = 0.004). Maternal rank had a strong effect on LRS (ß_Low maternal rank_ = −1.256 ± 0.232, p < 0.0001), and so did the mother’s death before the juvenile reached adulthood (ß_Mother died_ = −0.982± 0.300, p = 0.001).

**Figure 3.**
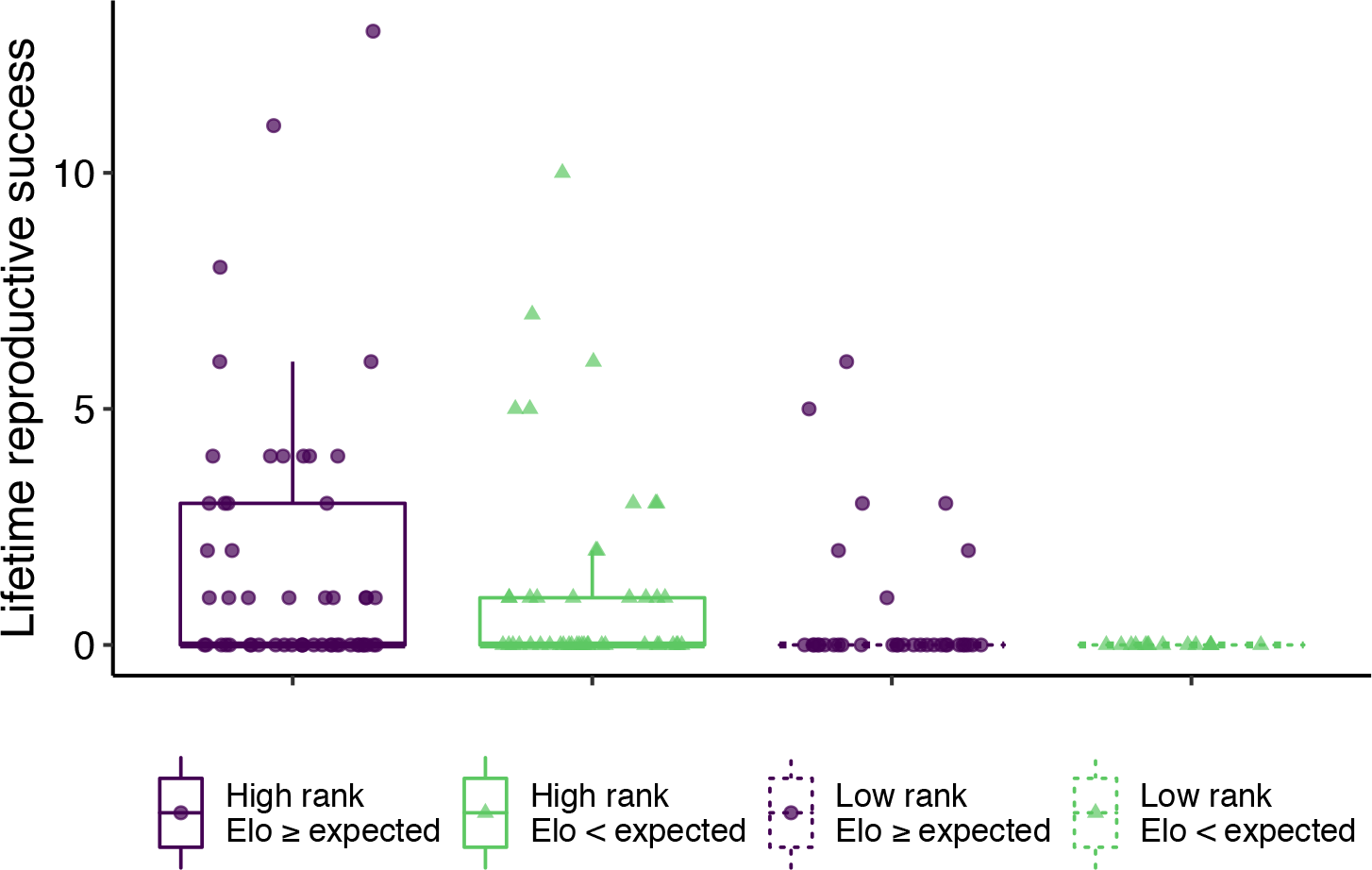
Lifetime reproductive success (LRS) as a function of both Elo-deviance at den independence and maternal rank. Juveniles with Elo-deviance < 0 showed reduced LRS, as did juveniles born to low ranking mothers. Death of the mother before the juvenile reached adulthood also predicted reduced LRS (not depicted).

## DISCUSSION

Our results reveal that, although rank acquisition follows a very predictable pattern of maternal rank inheritance with youngest ascendency in spotted hyenas (Figure 1), this process varies considerably among individuals, and this variation has profound consequences for both survival (Figure 2) and reproductive success (Figure 3). Individuals who tended to lose to their lower-born peers during the den dependent period (thus incurring an Elo-deviance below 0) suffered higher mortality and lower reproductive success than did those who won those fights.

Our results demonstrate that the ontogeny of dominance is related to fitness in ways that are not explained simply by the social status that juveniles attain as adults. In fact, we found that, depending on the measure of fitness, transient variation in the rank acquisition process can relate to fitness even more strongly than maternal rank (Figure 2). Here we found that the state of rank acquisition at den independence predicted survival and reproduction (Figures 2,3) but did not predict variation in the ranks attained as adults (Figure 1b). This result suggests that studies that focus on social status in adulthood overlook important potential rank-related influences on fitness that occur earlier during ontogenetic development.

How could transient variation in rank acquisition relate to fitness independent of adult rank? One interpretation is that variation in rank acquisition in juveniles is a source of early life hardship. Considerable evidence suggests that adverse conditions in early in life can have profound and long-lasting consequences [26]. Social defeat and social uncertainty in dominance relationships have been shown to incur costs [24,25,41]. Here, juveniles that were defeated by peers that they would eventually come to dominate showed reduced survival and impaired reproductive success, suggesting that social uncertainty coupled with social defeat could be a source of early life adversity in spotted hyenas. Our results are consistent with this suggestion. If we recode the three significant predictor variables from our fitness models (Above/below 0 deviance at den independence, High/low maternal rank, Mother alive/dead when juvenile reaches adulthood) into a single variable that counts the number of adverse conditions experienced by each juvenile, the number of early life adverse conditions significantly predicts increased mortality (hazard ratio = 1.522; 95% CI = [1.259, 1.840]; p < 0.0001; Figure 4). These results demonstrate that the adverse conditions studied here have cumulative effects, in that juveniles experiencing multiple adverse conditions suffer the additive combination of the consequences of each. In some species [26], multiple sources of early life adversity have compounding effects, in which the combination of sources of adversity have more severe consequences than the sum of the independent effects of each. We did not find any evidence for compounding effects here: the model with number of adverse conditions performed negligibly better than the original model that included each source of adversity as a separate fixed effect (AICc = 1.004), and a model including interactions between the adverse conditions performed more poorly than the model without interactions (∆AICc = 6.576).

**Figure 4.**
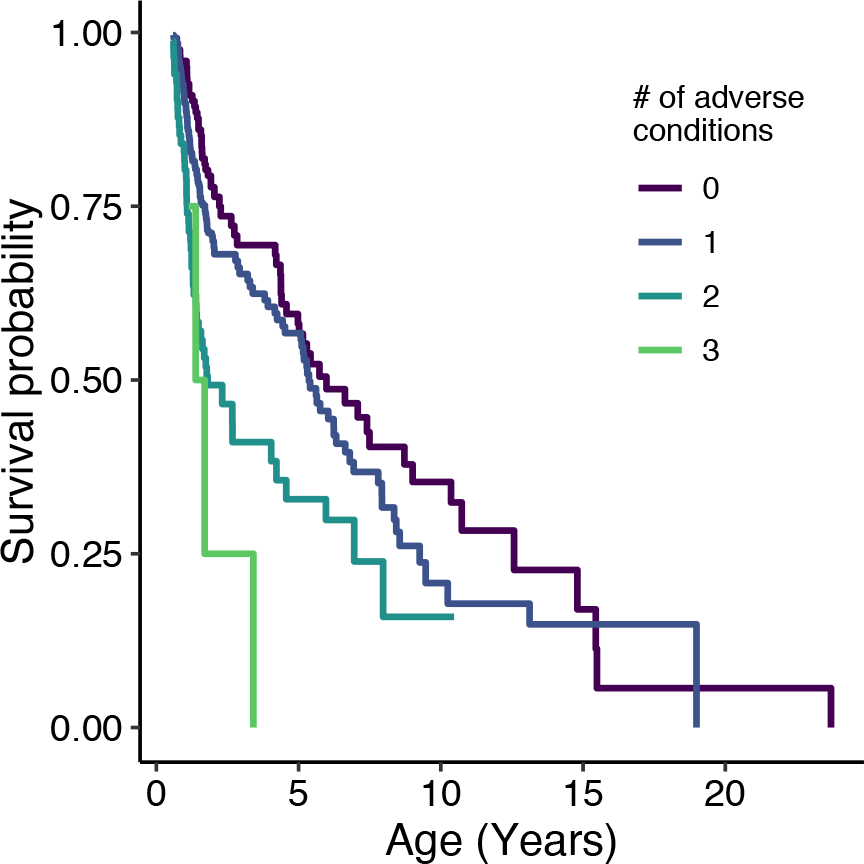
Survival probability as a function of the number of adverse conditions faced by juveniles during early life. The adverse conditions considered here were below Elo-deviance < 0 at den independence, low maternal rank, and death of mother before reaching adulthood.

The Elo-deviance method introduced here has proven to be a powerful tool for measuring deviation from a hypothesized pattern of contest outcomes. It’s ease of implementation, its customizability for addressing different questions, and its applicability with any hypothesis makes this a valuable new tool in studying animal dominance structures. To demonstrate how this method can be applied to ask a different question, in the Supplemental Materials we use the Elo-deviance method in a different way to investigate the timing of rank acquisition by juveniles.

Our work leaves open the question of what causes variation in rank acquisition. For example, variation in rank acquisition could be due to intrinsic differences between juveniles in quality or temperament. The fact that measures of rank acquisition calculated independently at different life-history stages were correlated is consistent with this conjecture. However, prior studies in spotted hyenas and other species with nepotistic societies suggest that mothers and other kin play an important role in the rank acquisition process, so the variation we observed here could also be sensitive to the behaviors of kin. For example, mothers may vary in their ability to support the process of rank acquisition of their juvenile offspring. If so, this may have important implications for variation across maternal lineages in the ability to sustain rank across generations. More generally, our work may provide a new piece to the puzzle of how maternal rank inheritance has evolved—if selection acts against those that deviate from the convention of maternal rank inheritance, then behavioral strategies may evolve to promote strict adherence to this convention and to enforce adherence by kin and other group-mates.

## Notes

https://github.com/straussed/rank_acquisition

## References

1. Holekamp KE, Strauss ED: Aggression and dominance: an interdisciplinary overview. Curr Opin Behav Sci 2016, 12:44–51.

2. Goymann W, Wingfield JC: Allostatic load, social status and stress hormones: The costs of social status matter. Anim Behav 2004, 67:591–602.

3. Strum SC: Agonistic dominance in male baboons: an alternative view. Int J Primatol 1982, 3:175–202.

4. Borries C, Sommer V, Srivastava A: Dominance, age, and reproductive success in free-ranging female hanuman langurs (Presbytis entellus). Int J Primatol 1991, 12:231–257.

5. Schülke O, Bhagavatula J, Vigilant L, Ostner J: Social bonds enhance reproductive success in male macaques. Curr Biol 2010, 20:2207–2210.

6. Silva LR, Lardy S, Ferreira AC, Rey B, Doutrelant C, Covas R: Females pay the oxidative cost of dominance in a highly social bird. Anim Behav 2018, 144:135–146.

7. East ML, Hofer H: Male spotted hyenas (Crocuta crocuta) queue for status in social groups dominated by females. Behav Ecol 2001, 12:558–68.

8. Haley MP, Deutsch CJ, Le Boeuf BJ: Size, dominance and copulatory success in male northern elephant seals, Mirounga angustirostris. Anim Behav 1994, 48:1249–1260.

9. Archie EA, Morrison TA, Foley CAH, Moss CJ, Alberts SC: Dominance rank relationships among wild female African elephants, Loxodonta africana. Anim Behav 2006, 71:117–127.

10. Tibbetts EA, Dale J: A socially enforced signal of quality in a paper wasp. Nature 2004, 432:218–222.

11. Barrette C, Vandal D: Social rank, dominance, antler size, and access to food in snow-bound wild woodland caribou. Behaviour 1986,

12. Wright E, Galbany J, McFarlin SC, Ndayishimiye E, Stoinski TS, Robbins MM: Male body size, dominance rank and strategic use of aggression in a group-living mammal. Anim Behav 2019, 151:87–102.

13. Strauss ED, Holekamp KE: Social alliances improve rank and fitness in convention-based societies. Proc Natl Acad Sci 2019, doi:10.1073/pnas.1810384116.

14. Vullioud C, Davidian E, Wachter B, Rousset F, Courtiol A, Höner OP: Social support drives female dominance in the spotted hyaena. Nat Ecol Evol 2019, 3:71–76.

15. Lea AJ, Learn NH, Theus MJ, Altmann J, Alberts SC: Complex sources of variance in female dominance rank in a nepotistic society. Anim Behav 2014, 94:87–99.

16. Dugatkin LA, Druen M: The social implications of winner and loser effects. Proc R Soc B Biol Sci 2004, 271:488–489.

17. Franz M, McLean E, Tung J, Altmann J, Alberts SC: Self-organizing dominance hierarchies in a wild primate population. Proc R Soc B Biol Sci 2015, 282:20151512.

18. Hiadlovská Z, Mikula O, Macholán M, Hamplová P, Vošlajerová Bímová B, Daniszová K: Shaking the myth: Body mass, aggression, steroid hormones, and social dominance in wild house mouse. Gen Comp Endocrinol 2015, 223:16–26.

19. Hass CC, Jenni DA: Structure and ontogeny of dominance relationships among bighorn rams. Can J Zool 1991, 69:471–476.

20. Holekamp KE, Smale L: Dominance acquisition during mammalian social development: the “inheritance” of maternal rank. Am Zool 1991, 31:306–317.

21. Tsai YJJ, Mann J: Dispersal, philopatry, and the role of fission-fusion dynamics in bottlenose dolphins. Mar Mammal Sci 2013, 29:261–279.

22. Stanton MA, Mann J: Early Social Networks Predict Survival in Wild Bottlenose Dolphins. PLoS One 2012, 7:1–6.

23. McDonald DB: Predicting fate from early connectivity in a social network. Proc Natl Acad Sci 2007, 104:10910–10914.

24. Vandeleest JJ, Beisner BA, Hannibal DL, Nathman AC, Capitanio JP, Hsieh F, Atwill ER, McCowan B: Decoupling social status and status certainty effects on health in macaques: a network approach. PeerJ 2016, 4:e2394.

25. Beaulieu M, Mboumba S, Willaume E, Kappeler PM, Charpentier MJE: The oxidative cost of unstable social dominance. J Exp Biol 2014, 217:2629–2632.

26. Tung J, Archie EA, Altmann J, Alberts SC: Cumulative early life adversity predicts longevity in wild baboons. Nat Commun 2016, 7:1–7.

27. Burton T, Metcalfe NB: Can environmental conditions experienced in early life influence future generations?Proc R Soc B Biol Sci 2014, 281:20140311.

28. Farine DR, Spencer KA, Boogert NJ: Early-Life Stress Triggers Juvenile Zebra Finches to Switch Social Learning Strategies. Curr Biol 2015, 25:2184–2188.

29. Smale L, Frank LG, Holekamp KE: Ontogeny of dominance in free-living spotted hyaenas: juvenile rank relations with adult females and immigrant males. Anim Behav 1993, 46:467–477.

30. Neumann C, Duboscq J, Dubuc C, Ginting A, Irwan AM, Agil M, Widdig A, Engelhardt A: Assessing dominance hierarchies: validation and advantages of progressive evaluation with Elo-rating. Anim Behav 2011, 82:911–921.

31. Elo AE: The rating of chessplayers, past and present. Arco Pub.; 1978.

32. Farine DR, Sanchez-Tojar A: aniDom: Inferring Dominance Hierarchies and Estimating Uncertainty. 2017,

33. Albers PCH, de Vries H: Elo-rating as a tool in the sequential estimation of dominance strengths. Animal 2001, 61:489–495.

34. Holekamp KE, Smale L: Ontogeny of dominance in free-living spotted hyaenas: juvenile rank relations with other immature individuals. Anim Behav 1993, 46:451–466.

35. Aureli F, Schaffner CM, Boesch C, Bearder SK, Call J, Chapman CA, Connor R, Fiore A Di, Dunbar RIM, Henzi SP, et al.: Fission-Fusion Dynamics. Curr Anthropol 2008, 49:627–654.

36. Altmann J: Observational study of behavior: sampling methods. Behaviour 1974, 49:227–266.

37. Strauss ED, Holekamp KE: Inferring longitudinal hierarchies: Framework and methods for studying the dynamics of dominance. J Anim Ecol 2019, 88:521–536.

38. Therneau TM: coxme: Mixed Effects Cox Models. R package version 2.2-10. 2018,

39. Strauss ED: DynaRankR: Inferring Longitudinal Dominance Hierarchies. 2019,

40. Bates D, Mächler M, Bolker B, Walker S: Fitting Linear Mixed-Effects Models Using lme4. J Stat Softw 2015, 67.

41. Sapolsky RM: The influence of social hierarchy on primate health. Science (80-) 2005, 308:648–652.

