## Supplemental Materials for "Juvenile rank acquisition influences fitness independent of adult rank"

#### 1. Spotted hyena life-history

Prior research has described how developing juvenile spotted hyenas pass through three important life-history stages before late adulthood. First, juveniles are typically born in litters of 1-2 at an isolated natal den, where they reside for the first 2-3 weeks of life. Births are rarely observed, so birthdates for cubs are estimated to within +/- 7 days based on the cubs' appearance when first observed [1].  Second, after 2-3 weeks, mothers move their offspring to a communal den to reside with all other juveniles within the clan until they are around 8-12 months old. During this den-dependent stage, juveniles rarely stray more than a few hundred meters from the shelter of den holes, and they regularly enter the den while resting or when threatened. Rank relationships among juvenile members of each cohort emerge while cubs live at the communal den. At the start of the communal den period, maternal rank has little influence on rank relationships among juvenile peers, but juvenile ranks closely match the maternal hierarchy by the time cubs become den-independent [2]. Third, juveniles achieve den-independence at around 8-12 months of age; as has been done before, we defined den-independence as the date on which a juvenile had been observed over 200m from the den on four consecutive occasions [3]. During the den-independent life-history stage, juveniles no longer reside at the den, but instead travel freely throughout the territory and associate in subgroups with related and unrelated group-mates. Weaning takes place during this den-independent period. After reaching reproductive maturity at 2 years old, males typically begin to disperse to new clans where they can be reproductively active, whereas females begin reproducing in their natal clans.

#### 2. Elo deviance at other life-history stages

In the main text, we assessed Elo deviance at den independence. To examine the state of rank acquisition over time, we also assessed Elo deviance at two later life-history stages. We calculated the state of rank acquisition at reproductive maturity (2 years old) as the Elo deviance calculated from interactions among den-independent juveniles (less than 2 years old) and the state of rank acquisition at the end of the first year of adulthood (3 years old) based on the interactions between these same individuals and all other adults. Importantly, these scores were not influenced by any interactions prior to den-independence because scores were 'reset' between life-history stages. At each life-history stage, we calculated Elo deviances for only those individuals who survived to the end of the period over which we calculated Elo deviance for that life-history stage and only those individuals who were observed engaging in aggressive interactions during this period.

Although Elo deviance at den independence (main text) was significantly correlated with Elo deviance at adulthood (Pearson's r = 0.193; 95% CI = [0.095, 0.288]; p = 0.0001, n = 385) and after the first year of adulthood (Pearson's r = 0.145; 95% CI = [0.011, 0.274]; p = 0.035, n = 213), models with Elo deviance assessed during these later life-history stages did not predict survival (Elo deviance < 0 at adulthood: n = 385; Hazard ratio = 1.042; 95% CI = [0.725, 1.500]; p = 0.82; Elo deviance < 0 after first year of adulthood: n = 215; Hazard ratio = 0.851; 95% CI = [0.509, 0.1.421]; p = 0.54). In models of LRS, Elo deviance class calculated at onset of adulthood (ß$_{\text{Elo deviance below 0}}$ = -0.310 ± 0.191, p = 0.104) and Elo deviance class calculated after the first year of adulthood (ß$_{\text{Elo deviance below 0}}$ = -0.238 ± 0.243, p = 0.164) were not significant predictors. In these models of survival and LRS at later life-history stages, covariates included were the number of interactions used to calculate Elo deviance during the different life-history stages, maternal rank (high or low), and a random effect of clan.

**3. Assessing the average timing of rank acquisition**

We used the Elo-deviance values to estimate the age (in months after birth) at which juveniles acquire their ranks based on maternal rank ‘inheritance.” We calculated Elo deviances for each observed individual in each month of life (from birth until death) using the individual’s interactions with all group-mates. We then summarized Elo deviances by month of age to investigate the variability in outcomes of dominance interactions at each month of age. Each individual had its Elo deviance calculated independently for each month of age (i.e., an individual’s score was ‘reset’ at each month of age).

We then measured the standard deviation of Elo deviances for all individuals at a given age. At ages where many individuals had contest outcomes that were not predicted by maternal rank, individuals had highly variable Elo deviances and thus that month of age had a large standard deviation in Elo deviances. At ages where contest outcomes of most individuals followed maternal rank, the standard deviation of Elo deviances for individuals at that month of age was closer to zero. To ensure that behavior at a given month of age was not unduly influenced by only a few individuals, months of age in which we had Elo deviances for fewer than 20 individuals were excluded from the analysis.

We expected the standard deviation of Elo deviances to decline during the early juvenile period up until some transition point at which most juveniles had fully acquired their maternal rank; after this transition point, we expected the standard deviation of Elo deviances to remain relatively constant across later months of age. To determine the month of age at which this transition takes place, we used piece-wise linear regression; we modeled the standard deviation of Elo deviances at each month of age as a function of age, and estimated a single break point using the bootstrap restarting algorithm implemented in the *segmented* R package [4,5].

Variation in Elo deviances binned by month of age declined steeply ($ß = -3.134 \pm 0.293$, $p < 0.0001$) until just after the first year of life (break-point = 12.97 months; Davie’s test $p < 0.0001$), after which deviance scores increased minimally over the remainder of life ($ß = 0.062 \pm 0.029$, $p = 0.035$) (Figure S1). Examination of individual Elo deviance scores assessed relative to their peers at the three different life-history stages shows a similar pattern. The standard deviation of Elo deviance at den independence (sd = 62.171, n = 465) was around double the standard deviation of Elo deviance at onset of adulthood (sd = 32.723, n = 385) and after the first year of adulthood (sd = 32.442, n = 213).  Our results confirmed our expectations, and produced an estimate of completion of rank acquisition (13.05 months) that was slightly earlier than the rough estimate of 18 months from prior work [6].

These results, in conjunction with the results presented in the main text, demonstrate the flexibility of the Elo deviance method and it's utility in addressing diverse questions. By tailoring the time-frame at which deviance scores are assessed (den-dependent life-history stage in main text vs. monthly over the entire lifespan here), the types of interactions used to calculate the Elo deviance (only interactions with other juveniles in main text vs. interactions with any clan member here), and the hypothesis being considered (maternal rank 'inheritance' in both cases), this method can be used to ask a variety of interesting questions about dominance-related behavior.

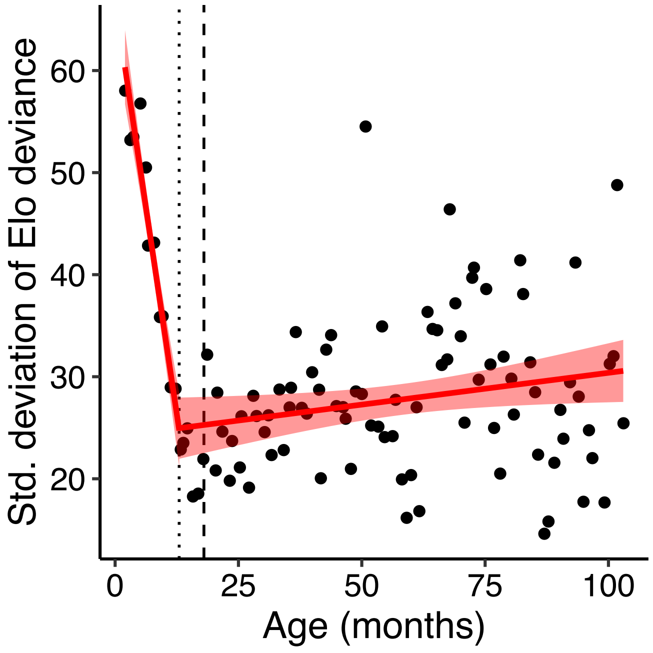

**Figure S1**. The timing of the development of juvenile social status. Piecewise linear regression revealed that juvenile Elo deviances were highly variable up until 12.97 months (dotted line), after which their variability was comparable to that of adults. This estimate of the timing of the establishment of social status resembles the 18 months (dashed line) described previously [Smale 1993].

### 4. Parameterization of Elo-rating

The most important parameter in the Elo-rating method is K, which is a constant that influences the magnitude of changes in scores after each interaction. This constant is weighted by the expected probability of the outcome, such that unexpected outcomes result in changes closer to K and expected outcomes result in changes closer to 0. We used K = 20 for the analyses in the main text, but here we provide the plots from analyses with K = 100. Varying this parameter had no effect on the conclusions of our study.

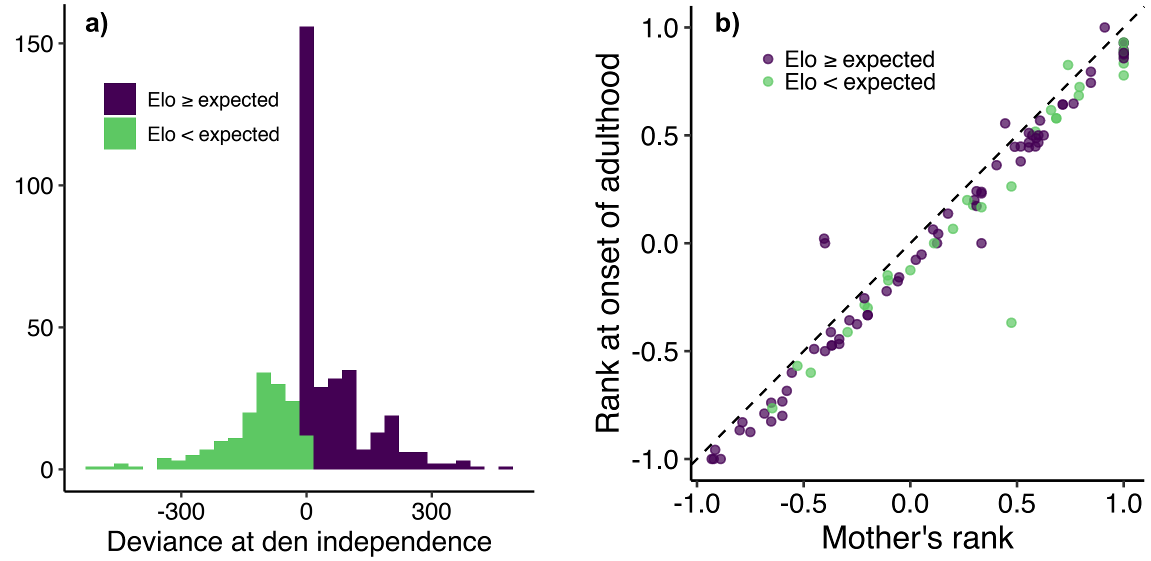


**Figure S2**. (a) Histogram of Elo deviance at den independence. (b) The relationship between the juvenile's mother's rank and the juvenile's rank at onset of adulthood (2 years of age). According to maternal rank inheritance, points should lie directly below the dashed line (denoting where mother's rank and juvenile's rank are exactly equal). In this study, 77.5% of juveniles acquired the exact rank predicted by maternal rank inheritance. Elo deviance at den independence (color) did not affect the rank attained by the onset of adulthood. Taken together, these plots show transient variability in rank acquisition at the end of den-dependent life-history stage that doesn't manifest in differences in rank at adulthood. Elo deviance was calculated here with the Elo-rating parameter K = 100 rather than K = 20 in the main text (Figure 1).

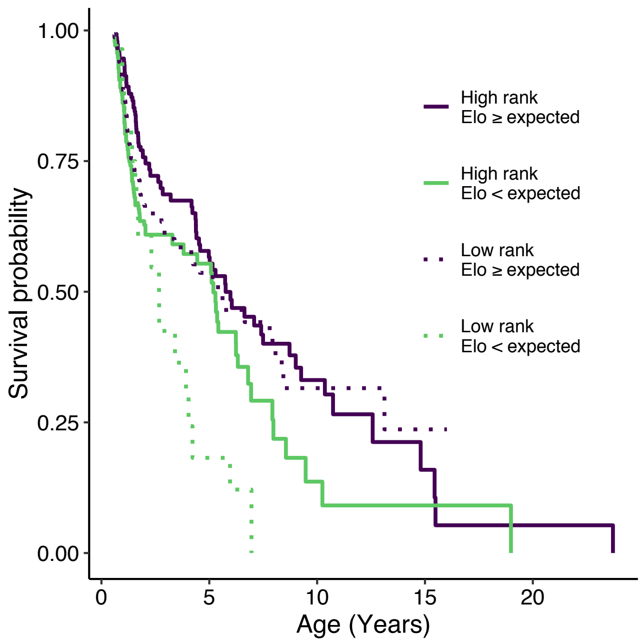


**Figure S3**. Survival probability as a function of Elo deviance at den independence and maternal rank. Juveniles with below Elo deviance < 0 showed reduced survival. Death of the mother before the juvenile reached adulthood also predicted reduced survival, but being born to a low-ranking mother did not predict survival after controlling for the other variables in the model. Elo deviance was calculated here with the Elo-rating parameter K = 100 rather than K = 20 in the main text (Figure 2).

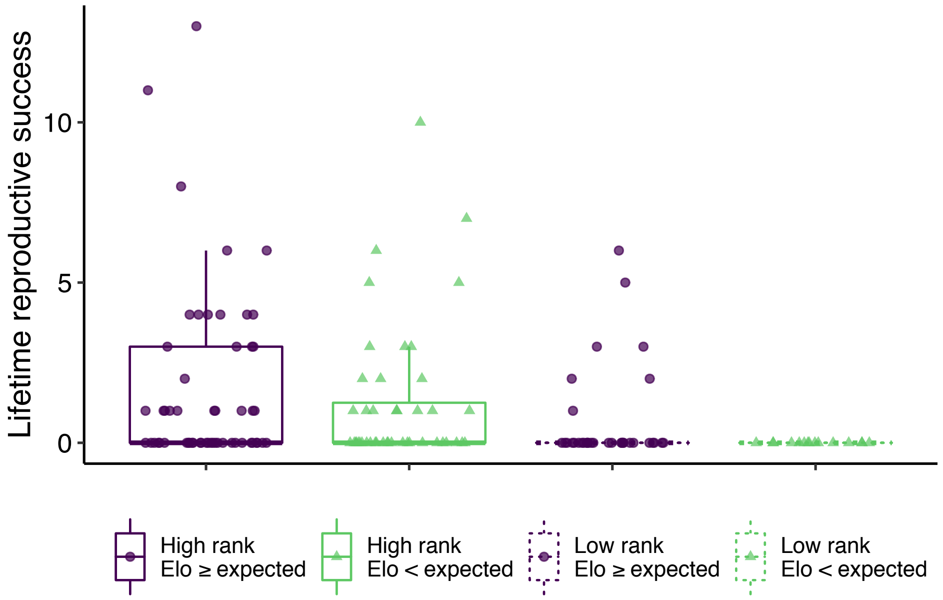


**Figure S4**. Lifetime reproductive success (LRS) as a function of Elo deviance at den independence and maternal rank. Juveniles with Elo deviance < 0 showed reduced LRS, as did juveniles born to low ranking mothers. Death of the mother before the juvenile reached adulthood also predicted reduced LRS (not depicted). Elo deviance was calculated here with the Elo-rating parameter K = 100 rather than K = 20 in the main text (Figure 3).

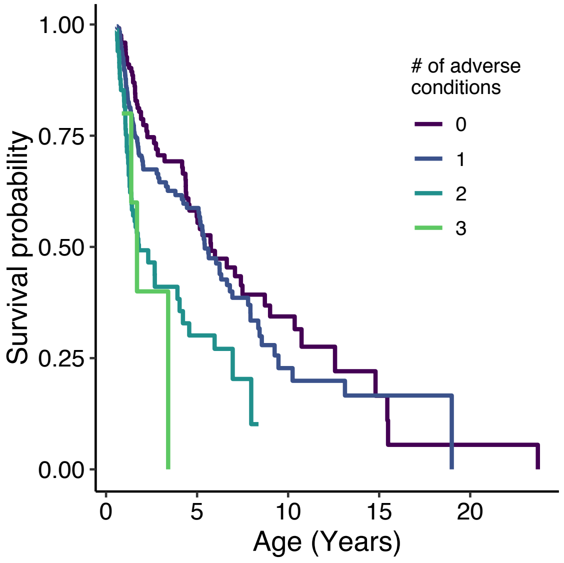


**Figure S5**. Survival probability as a function of the number of adverse conditions faced by juveniles during early life. The adverse conditions considered here were below Elo deviance < 0 at den independence, low maternal rank, and death of mother before reaching adulthood. Elo deviance was calculated here with the Elo-rating parameter K = 100 rather than K = 20 in the main text (Figure 4).

### 5. References


1. Holekamp KE, Smale L, Szykman M: **Rank and reproduction in the female spotted hyaena**. *J Reprod Fertil* 1996, **108**:229–237.
2. Holekamp KE, Smale L: **Ontogeny of dominance in free-living spotted hyaenas: juvenile rank relations with other immature individuals**. *Anim Behav* 1993, **46**:451–466.
3. Boydston EE, Kapheim KM, Van Horn RC, Smale L, Holekamp KE: **Sexually dimorphic patterns of space use throughout ontogeny in the spotted hyena (Crocuta crocuta)**. *J Zool* 2005, **267**:271.
4. Muggeo VR: **segmented: an R Package to Fit Regression Models with Broken-Line Relationships**. *R News* 2008,
5. Muggeo VMR: **Estimating regression models with unknown break-points**. *Stat Med* 2003, **22**:3055–3071.
6. Smale L, Frank LG, Holekamp KE: **Ontogeny of dominance in free-living spotted hyaenas: juvenile rank relations with adult females and immigrant males**. *Anim Behav* 1993, **46**:467–477.
